# Deletion of the ribosomal protein gene *rpmJ* activates *zntA* transcription through a translation-dependent mechanism in *Escherichia coli*

**DOI:** 10.64898/2026.08.06.743257

**Authors:** Riko Shirakawa, Kazuya Ishikawa, Kazuyuki Furuta, Chikara Kaito

## Abstract

Bacteria tightly regulate intracellular zinc homeostasis by coordinating zinc uptake and efflux systems. We previously showed that deletion of the ribosomal protein gene *rpmJ* confers zinc resistance in *Escherichia coli* in a manner dependent on the zinc efflux transporter *zntA*. Here, we analyzed the effect of *rpmJ* deficiency on *zntA* expression. Under zinc excess conditions, *zntA* mRNA levels were markedly higher in the *rpmJ* mutant than in the wild-type strain. Enhanced *zntA* expression required the native *zntA* promoter, the native Shine–Dalgarno sequence, and the N-terminal coding region of *zntA*, indicating that translation initiated from the native Shine–Dalgarno sequence and extending through the N-terminal coding region is required for enhanced transcription initiation from the native *zntA* promoter. Furthermore, ectopic expression of *ykgO*, a paralog of *rpmJ* that is known to replace RpmJ on the ribosome under zinc-limited conditions, abolished the increased *zntA* expression and zinc resistance conferred by *rpmJ* deletion. Collectively, these findings suggest that ribosomes lacking RpmJ or YkgO promote transcription initiation from the native *zntA* promoter through translation of *zntA* mRNA.

**IMPORTANCE:** Bacteria must carefully control the amount of zinc inside their cells. Too little zinc prevents essential cellular processes, whereas too much zinc is toxic. We found that removing a small ribosomal protein called RpmJ allows *Escherichia coli* to survive high zinc levels by increasing production of the zinc exporter *zntA*. Surprisingly, this increase depends not only on the *zntA* promoter but also on translation of the beginning of the *zntA* coding region. Our findings suggest that changes in ribosome composition can stimulate gene transcription through early translation of the same messenger RNA, revealing a previously unrecognized mechanism linking translation and transcription during bacterial adaptation to zinc stress.

## INTRODUCTION

Zinc is an essential trace element that serves as a catalytic cofactor and structural component of numerous proteins and is indispensable for cellular functions in organisms ranging from bacteria to humans. Excess intracellular zinc, however, is toxic because it disrupts iron–sulfur clusters (1) and promotes mismetalation of metalloproteins by competing with other essential metal ions such as manganese (2, 3). To maintain zinc homeostasis, bacteria tightly regulate zinc uptake and efflux systems. In *Escherichia coli*, the high-affinity zinc uptake system *znuABC* is induced under zinc-deficient conditions (4), whereas the zinc efflux transporter *zntA* is induced under zinc-excess conditions (5, 6), thereby maintaining intracellular zinc concentrations within a narrow physiological range. In addition to these transport systems, exposure to excess zinc alters the expression of numerous genes that are not directly involved in zinc transport (7, 8). These observations suggest that bacterial adaptation to zinc stress involves regulatory mechanisms beyond zinc uptake and efflux, although the molecular basis of these responses remains largely unknown.

The *E. coli* ribosome consists of 54 ribosomal proteins. Among them, RpmJ (L36), the smallest protein in the 50S subunit (9), contains a zinc ribbon motif that coordinates zinc ions (10, 11). RpmJ has a paralog, YkgO, which lacks a zinc-binding motif and occupies the same position in the ribosome as RpmJ (12). Likewise, the L31 ribosomal protein RpmE has a paralog, YkgM, whose product also lacks a zinc-binding motif. Under zinc-replete conditions, expression of *ykgO* and *ykgM* is repressed by the zinc-responsive transcriptional regulator Zur (13, 14). Under zinc-deficient conditions, this repression is relieved, allowing YkgO and YkgM to replace RpmJ and RpmE, respectively, in the ribosome while maintaining translational capacity (15). During this replacement, RpmJ and RpmE released from the ribosome are thought to liberate zinc coordinated by their zinc-binding motifs, thereby increasing the intracellular pool of available zinc. Thus, the RpmJ–YkgO and RpmE–YkgM exchange systems are considered to function as a zinc-sparing mechanism that uses the ribosome as a dynamic zinc reservoir in response to zinc limitation. This adaptive strategy was first identified in *Bacillus subtilis*, where replacement of the zinc-binding ribosomal protein RpmE by its paralog YtiA was proposed to mobilize ribosome-associated zinc during zinc starvation (16).

We previously identified an *rpmJ* deletion mutant as a novel zinc-resistant strain through screening of the *E. coli* Keio collection (17). We further demonstrated that intracellular zinc levels are reduced in the *rpmJ* mutant and that deletion of the zinc efflux transporter gene *zntA* abolishes its zinc-resistant phenotype, suggesting that loss of *rpmJ* enhances ZntA-mediated zinc efflux. However, whether *rpmJ* deficiency directly affects *zntA* expression and, if so, how such regulation is achieved remained unknown.

In the present study, we investigated the effect of *rpmJ* deficiency on *zntA* expression and elucidated the molecular mechanism underlying its regulation. We show that deletion of *rpmJ* increases *zntA* transcription under zinc-excess conditions and that this induction requires translation of the N-terminal *zntA* coding region. Furthermore, because expression of the *rpmJ* paralog *ykgO* complemented the zinc-resistant phenotype of the *rpmJ* mutant, our findings indicate that zinc resistance results from altered ribosome composition caused by the absence of RpmJ or YkgO, rather than from the zinc-binding activity of RpmJ itself.

## RESULTS

### The *rpmJ*-deficient strain exhibits zinc resistance in a *zntR*–*zntA*-dependent manner

We previously showed that deletion of the zinc efflux transporter gene *zntA* abolishes the zinc-resistant phenotype of the *rpmJ*-deficient strain (17). To determine whether zinc uptake systems also contribute to this phenotype, we examined zinc resistance in mutants lacking the zinc uptake transporters *znuB*, a component of the high-affinity zinc transporter ZnuABC, or *zupT*, a ZIP family transporter (18). Deletion of either *znuB* or *zupT* did not affect the zinc resistance conferred by *rpmJ* deficiency (Fig. 1A). Consistent with our previous findings, *rpmJ* deficiency also conferred zinc resistance in the absence of the zinc efflux transporter *zitB* (19), whereas deletion of *zntA* completely abolished this phenotype (Fig. 1A). These results indicate that zinc resistance conferred by *rpmJ* deficiency specifically requires *zntA* but not the zinc uptake transporters *znuB* and *zupT* or the zinc efflux transporter *zitB*.

**Figure. 1.**
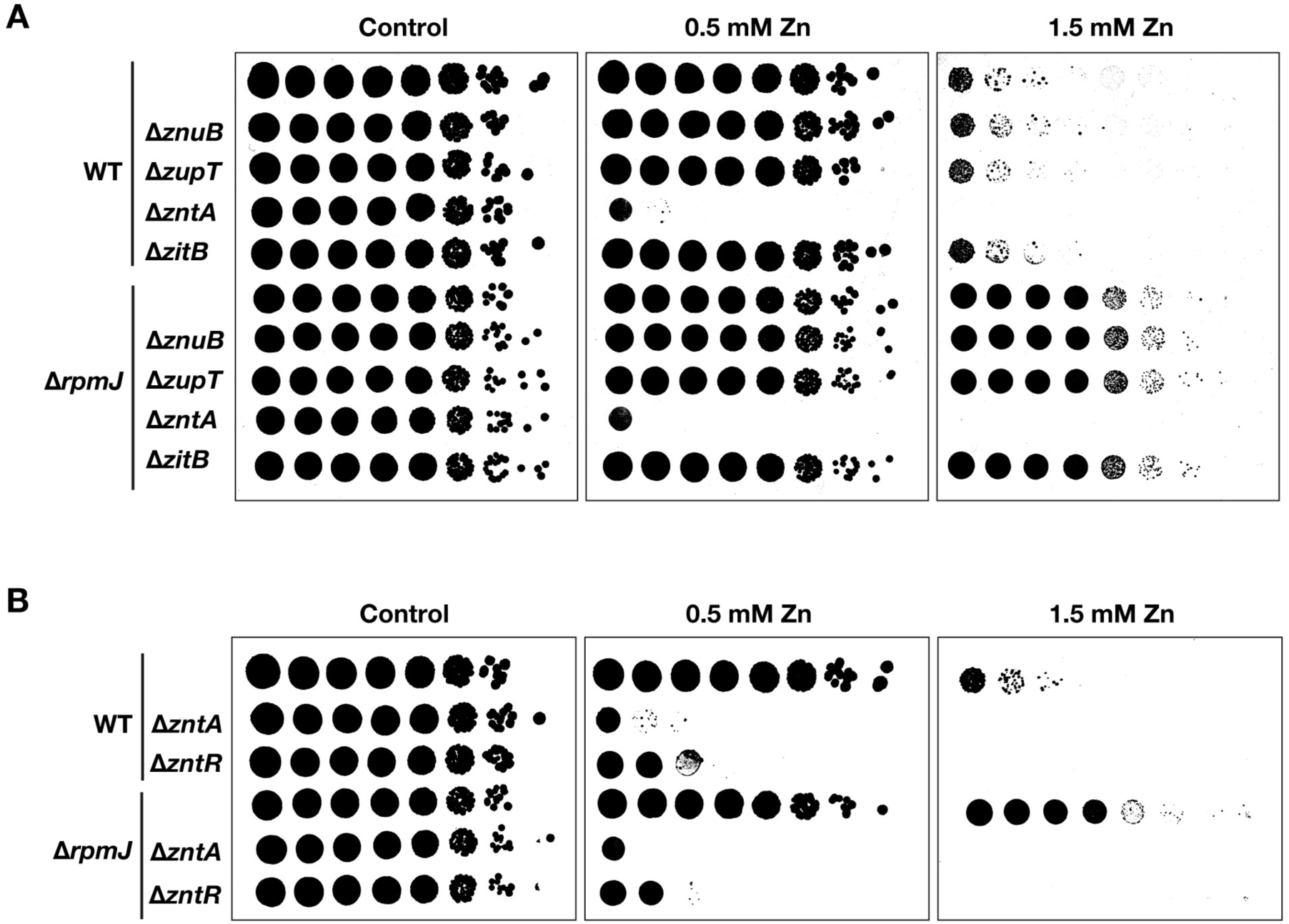
The *rpmJ*-deficient strain exhibits zinc resistance in a manner dependent on the zinc efflux transporter *zntA* and its transcriptional activator *zntR*. **(A)** Identification of factors required for the zinc resistance of the Δ*rpmJ* strain. Ten-fold serial dilutions of overnight cultures of the *E. coli* wild-type strain (BW25113), strains lacking zinc uptake or efflux factors, and strains lacking these same factors in the Δ*rpmJ* background were spotted onto LB agar plates containing 0.5 or 1.5 mM Zn(zinc) at a final concentration, or onto plain LB agar plates, and incubated for approximately 20 hours. Images are shown after binarization. **(B)** Involvement of the zinc efflux pathway in the zinc resistance of the Δ*rpmJ* strain. Ten-fold serial dilutions of overnight cultures of the *E. coli* wild-type strain, strains lacking *zntA* or its transcriptional activator *zntR*, and strains lacking these same factors in the Δ*rpmJ* background were spotted onto LB agar plates containing 0.5 or 1.5 mM Zn at a final concentration, or onto plain LB agar plates, and incubated for approximately 20 hours. Images are shown after binarization.

Expression of *zntA* is activated by the zinc-responsive transcriptional regulator ZntR (20, 21). We therefore examined whether *zntR* is also required for zinc resistance in the *rpmJ*-deficient strain. Deletion of *zntR* completely abolished the zinc-resistant phenotype of the *rpmJ* mutant (Fig. 1B). Together, these results demonstrate that zinc resistance conferred by *rpmJ* deficiency depends on the *zntR*–*zntA* regulatory pathway.

### Under zinc-excess conditions, *rpmJ* deficiency increases *zntA* expression without altering ZntR expression

Because zinc resistance conferred by *rpmJ* deficiency required both *zntA* and *zntR*, we examined whether loss of *rpmJ* affects the expression of either gene. Western blot analysis showed that ZntA protein levels increased in both the wild-type and Δ*rpmJ* strains at 0.5 and 1.0 mM Zn, with no significant difference between the strains (Fig. 2A). As the zinc concentration increased from 1.5 to 2.5 mM, ZntA protein levels declined in the wild-type strain but remained elevated in the Δ*rpmJ* strain. Consequently, at 2.5 mM Zn, ZntA protein levels in the Δ*rpmJ* strain were more than sixfold higher than those in the wild-type strain (Fig. 2A).

**Figure. 2.**
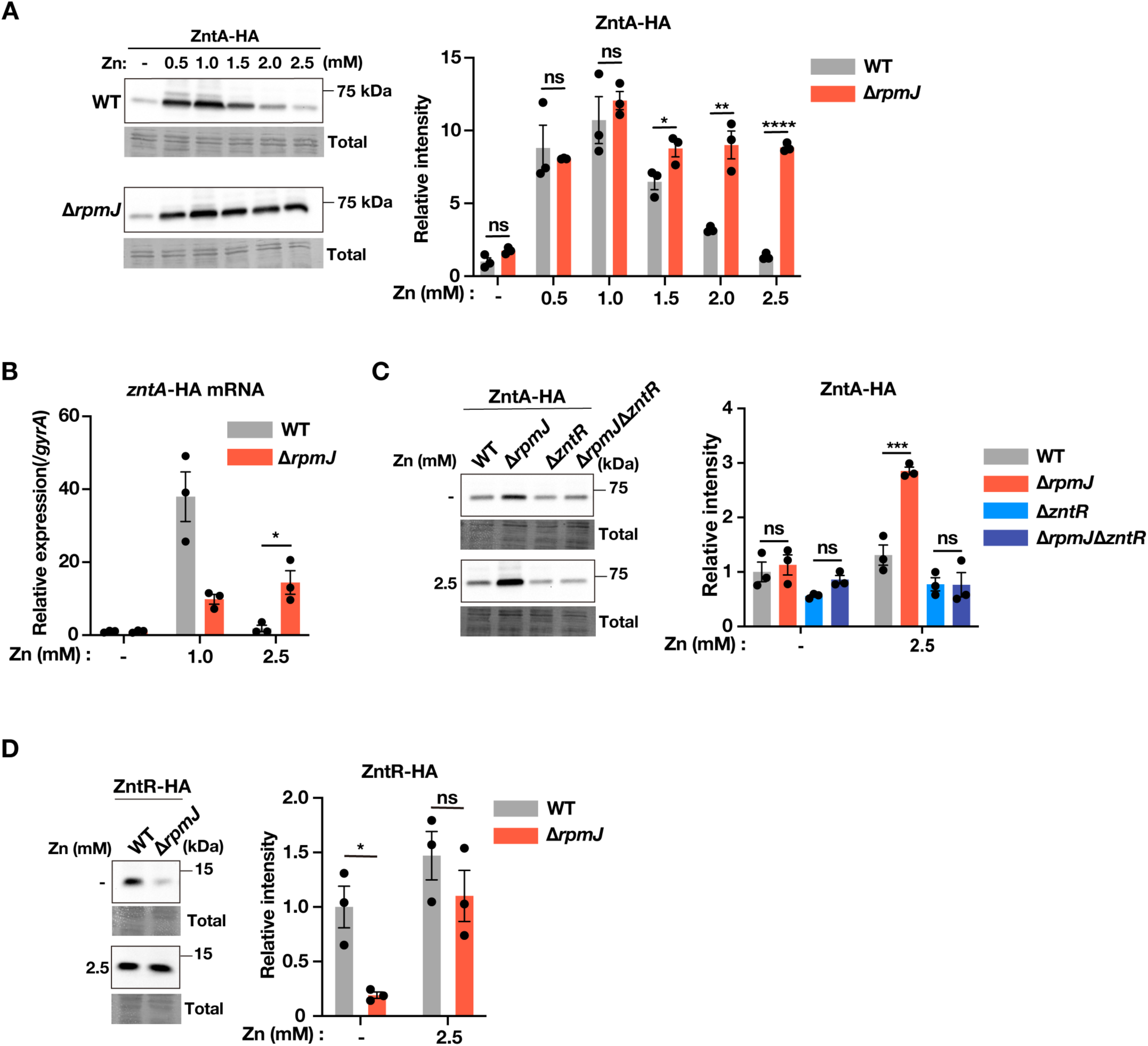
Under zinc-excess conditions, the Δ*rpmJ* strain exhibits increased *zntA* expression without increased ZntR expression. **(A)** ZntA-HA expression in the Δ*rpmJ* strain. pMW118-zntA-HA was introduced into the wild-type and Δ*rpmJ* strains. Cells were cultured for 2 hours, followed by the addition of zinc to final concentrations of 0.5, 1.0, 1.5, 2.0, or 2.5 mM and incubation for an additional 1 hour. Control cultures received no zinc. Cell lysates were analyzed by western blotting using an anti-HA antibody. Total protein levels were assessed by CBB staining of the membrane after transfer. Data are presented as mean ± S.E.M. (*n* = 3, \**p*< 0.05, \*\**p* < 0.01, \*\*\*\**p* < 0.0001; unpaired *t*-test). **(B)** *zntA*-HA mRNA expression in the Δ*rpmJ* strain. Cells were cultured as described in (A), and total RNA was analyzed by RT-qPCR. Transcript levels were normalized to *gyrA*. Data are presented as mean ± S.E.M. (*n* = 3, *\*p* < 0.05; unpaired *t*-test). **(C)** ZntA-HA expression in the Δ*zntR* and Δ *rpmJ* Δ *zntR* strains. Cells were cultured and analyzed as described in (A). Total protein levels were assessed by CBB staining of the membrane after transfer. Data are presented as mean ± S.E.M. (*n* = 3, \*\*\**p* < 0.001; Tukey’s multiple comparisons test). **(D)** ZntR-HA expression in the Δ*rpmJ* strain. pMW118-zntR-HA was introduced into the wild-type and Δ*rpmJ* strains. Cells were cultured and analyzed as described in (A). Total protein levels were assessed by CBB staining of the membrane after transfer. Data are presented as mean ± S.E.M. (*n* = 3, \**p* < 0.05; unpaired *t*-test).

We next quantified *zntA* mRNA levels. *zntA* mRNA levels in the Δ*rpmJ* strain were lower than those in the wild-type strain at 1.0 mM Zn but were increased more than sevenfold at 2.5 mM Zn (Fig. 2B), consistent with the increase in ZntA protein levels observed at this concentration. These results indicate that *rpmJ* deficiency enhances *zntA* expression specifically under zinc-excess conditions.

We next examined whether the elevated *zntA* expression required *zntR*. The increase in ZntA protein expression observed in the Δ*rpmJ* strain at 2.5 mM Zn was completely abolished in the Δ*zntR* background (Fig. 2C). We therefore examined whether *rpmJ* deficiency affects ZntR expression. Western blot analysis showed that ZntR protein levels were comparable between the wild-type and Δ*rpmJ* strains at 2.5 mM Zn (Fig. 2D). Together, these results indicate that the enhanced *zntA* expression caused by *rpmJ* deficiency requires *zntR* but is not attributable to increased ZntR expression.

### The *zntA* coding region is required for enhanced transcription initiation in the *rpmJ*-deficient strain

To determine whether increased transcription initiation contributes to the elevated *zntA* mRNA levels observed in the *rpmJ*-deficient strain under zinc-excess conditions, we first examined *zntA* promoter activity using a reporter containing the native *zntA* promoter and Shine–Dalgarno (SD) sequence fused to sfGFP (Fig. 3A). At 1.0 mM Zn, reporter activity was lower in the Δ*rpmJ* strain than in the wild-type strain (Fig. 3B). At 2.5 mM Zn, however, reporter activity was comparable between the two strains (Fig. 3B), indicating that the *zntA* promoter and SD sequence alone are insufficient to account for the increased *zntA* expression observed in the Δ*rpmJ* strain.

**Figure. 3.**
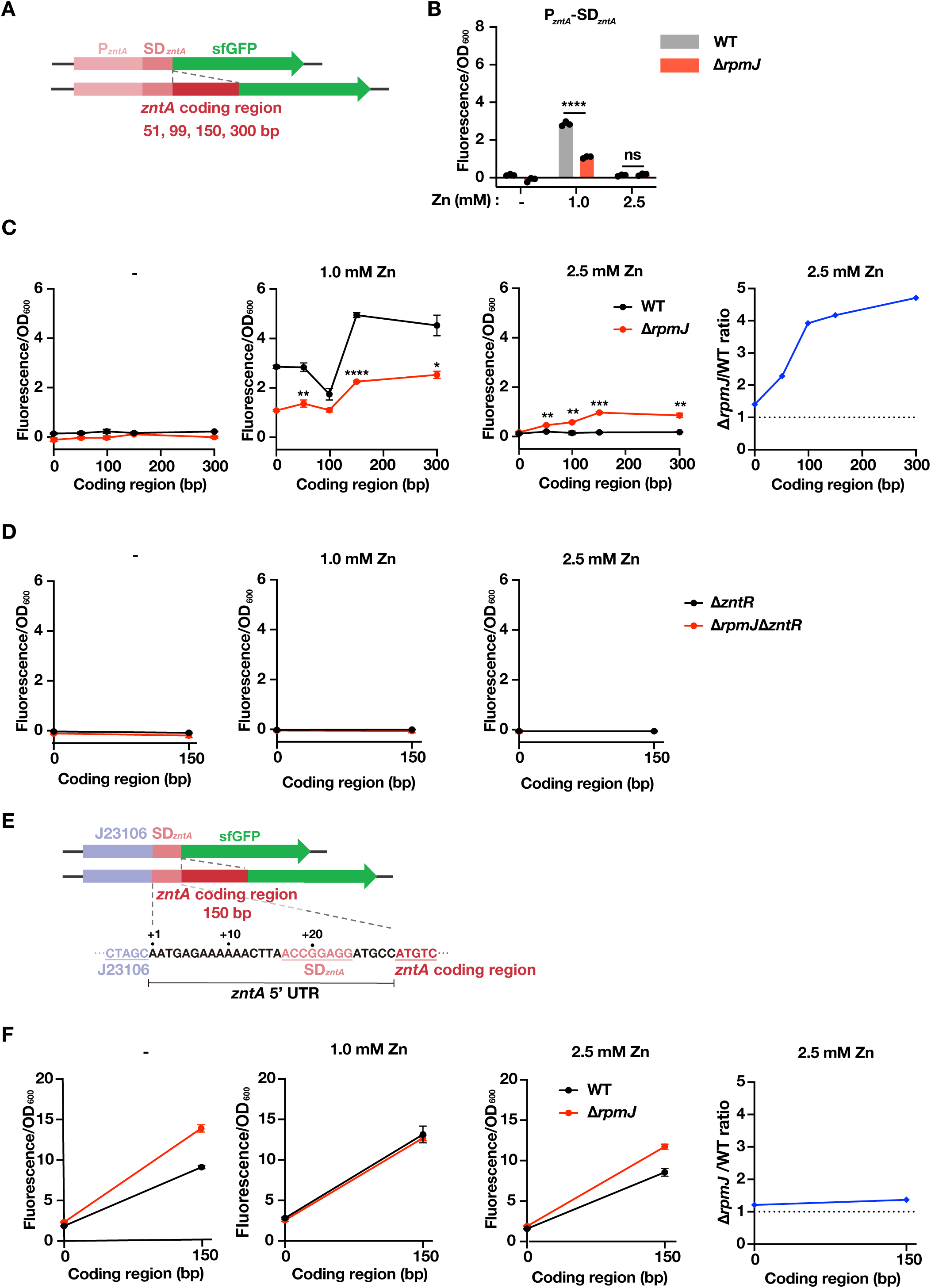
The *zntA* coding region is required for enhanced *zntA* expression in the Δ*rpmJ* strain under zinc-excess conditions. (**A**) Reporter constructs. The region extending from the *zntA* promoter to the Shine–Dalgarno sequence (P*zntA*-SD*zntA*) or constructs containing the first 51, 99, 150, or 300 bp of the *zntA* coding region downstream of the native promoter (P*zntA*-SD*zntA*-*zntA* coding region) were fused upstream of *sfGFP*. (**B**) Reporter activity of the P*zntA*-SD*zntA* construct. pMW118-P*zntA*-SD*zntA*-sfGFP was introduced into the wild-type and Δ*rpmJ* strains. Cells were cultured as described in Fig. 2A, and reporter activity was calculated from fluorescence intensity normalized to OD600. Data are presented as mean ± S.E.M. (*n* = 3, \*\*\*\**p* < 0.0001; unpaired *t*-test). (**C**) Reporter activity of constructs containing the *zntA* coding region. Reporter plasmids containing 51, 99, 150, or 300 bp of the *zntA* coding region were introduced into the wild-type and Δ*rpmJ* strains, and reporter activity was determined as described in (B). Data are presented as mean ± S.E.M. (*n* = 3, \**p*< 0.05, \*\**p* < 0.01, \*\*\**p* < 0.001, \*\*\*\**p* < 0.0001; unpaired *t*-test at each length). The graph on the right shows the Δ*rpmJ*/WT ratio at 2.5 mM Zn. (**D**) Requirement of ZntR for enhanced reporter activity. The reporter construct containing the first 150 bp of the *zntA* coding region was introduced into the Δ*zntR* and Δ*rpmJ* Δ*zntR* strains, and reporter activity was determined as described in (B). (n=3) (**E**) Reporter construct in which the native *zntA* promoter was replaced with the constitutive promoter J23106. The 5′-untranslated region from the transcription start site (+1) to the start codon was retained. (**F**) Reporter activity of the J23106 promoter constructs. Reporter plasmids carrying J23106 with or without the first 150 bp of the *zntA* coding region were introduced into the wild-type and Δ*rpmJ* strains, and reporter activity was determined as described in (B). (n=3) The graph on the right shows the Δ*rpmJ*/WT ratio at 2.5 mM Zn.

We next generated a series of reporter plasmids in which progressively longer fragments of the *zntA* coding region were inserted between the native *zntA* promoter–SD sequence and sfGFP (Fig. 3A). At 1.0 mM Zn, insertion of either 150 bp or 300 bp of the *zntA* coding region increased reporter activity in both strains, although activity remained lower in the Δ*rpmJ* strain than in the wild-type strain (Fig. 3C). In contrast, at 2.5 mM Zn, reporter activity was barely detectable for all constructs in the wild-type strain, whereas it increased in proportion to the length of the inserted *zntA* coding region in the Δ*rpmJ* strain, reaching a maximum at 150 bp. (Fig. 3C). Reporter activity of the constructs containing 150 bp or 300 bp of the coding region was approximately four- to fivefold higher in the Δ*rpmJ* strain than in the wild-type strain (Fig. 3C, right).

Furthermore, deletion of *zntR* abolished the activity of both the promoter–SD reporter and the reporter containing an additional 150 bp of the *zntA* coding region in both the wild-type and Δ*rpmJ* strains (Fig. 3D). These results indicate that enhanced *zntA* transcription in the Δ*rpmJ* strain under 2.5 mM Zn conditions requires both *zntR* and the *zntA* coding region.

To further determine whether the increased *zntA* expression results from enhanced transcription initiation at the native promoter, we replaced the *zntA* promoter with the constitutive promoter J23106 (BBa_J23106) while preserving the native transcription start site and 5’-untranslated region (5’ UTR) (Fig. 3E). Under these conditions, reporter activity was comparable between the wild-type and Δ*rpmJ* strains regardless of the presence of the 150-bp *zntA* coding region or the zinc concentration (Fig. 3F). At 2.5 mM Zn, the reporter containing the 150-bp coding region showed only a 1.4-fold increase in the Δ*rpmJ* strain relative to the wild type (Fig. 3F, right). These findings indicate that *rpmJ* deficiency enhances transcription initiation from the native *zntA* promoter and that this regulation requires the *zntA* coding region.

### Translation initiation at the native *zntA* Shine–Dalgarno sequence is required for enhanced *zntA* transcription in the *rpmJ*-deficient strain

Because the *zntA* coding region was required for enhanced transcription initiation from the native *zntA* promoter in the Δ*rpmJ* strain under excess zinc conditions, we next examined whether translation initiation of *zntA* is required for this regulation. To inhibit translation initiation, the *zntA* start codon was replaced with a stop codon (Fig. 4A). Under 2.5 mM Zn conditions, the increase in *zntA* mRNA levels observed in the Δ*rpmJ* strain was attenuated by the stop codon mutation (Fig. 4B). The Δ*rpmJ*/WT ratio decreased from approximately fivefold for the wild-type construct to approximately threefold for the stop-codon mutant (Fig. 4B, right). These results indicate that translation initiation of *zntA* is required for the increase in *zntA* mRNA levels in the *rpmJ*-deficient strain under zinc-excess conditions.

**Figure. 4.**
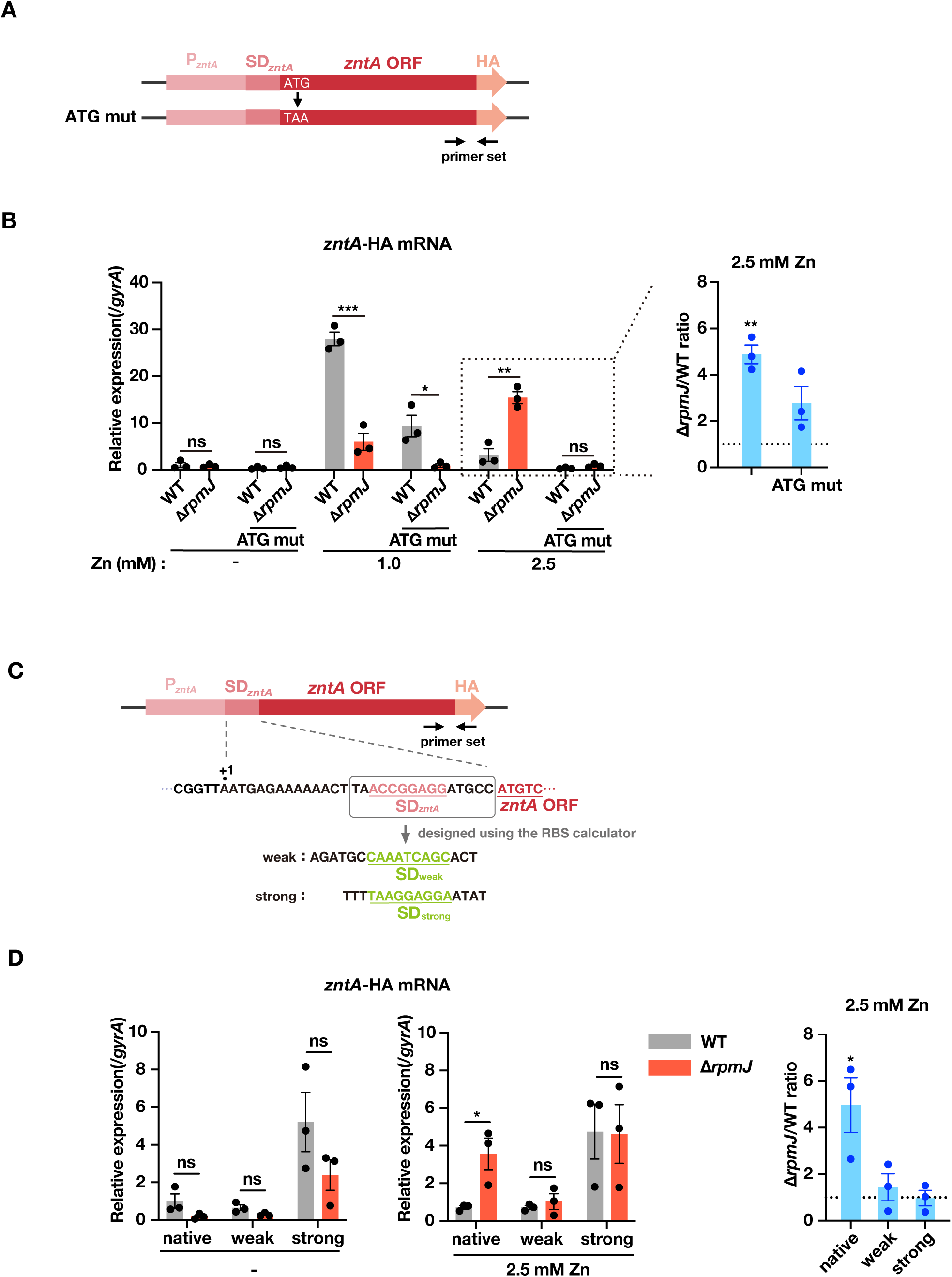
Translation initiation of *zntA* is required for enhanced *zntA* mRNA expression in the Δ*rpmJ* strain under zinc-excess conditions. **(A)** Construct carrying a mutation in the *zntA* start codon. The native *zntA* start codon (ATG) was replaced with the stop codon TAA. **(B)** Effect of the *zntA* start codon mutation on *zntA*-HA mRNA expression. pMW118-zntA-HA or pMW118-zntA-HA (ATG mutant) was introduced into the wild-type and Δ*rpmJ* strains. Cells were cultured and analyzed as described in Fig. 2B. Data are presented as mean ± S.E.M. (*n* = 3, *\*p* < 0.05, \*\**p* < 0.01, \*\*\**p* < 0.001; unpaired *t*-test). The graph on the right shows the Δ*rpmJ*/WT ratio at 2.5 mM Zn (\*\**p* < 0.01; unpaired *t*-test). **(C)** Constructs carrying mutations in the *zntA* Shine–Dalgarno sequence. Variants with weaker or stronger predicted translation initiation strength than the native Shine–Dalgarno sequence were designed using the RBS Calculator. **(D)** Effect of Shine–Dalgarno sequence mutations on *zntA*-HA mRNA expression. pMW118-zntA-HA, pMW118-zntA-HA (SDweak), or pMW118-zntA-HA (SDstrong) was introduced into the wild-type and Δ*rpmJ* strains. Cells were cultured and analyzed as described in Fig. 2B. Data are presented as mean ± S.E.M. (*n* = 3, \**p* < 0.05; unpaired *t*-test). The graph on the right compares the wild-type and Δ*rpmJ* strains for each construct (\**p* < 0.05; unpaired *t*-test).

We next examined whether translation initiation rate is involved in this regulation by constructing reporter variants carrying mutations in the *zntA* Shine–Dalgarno (SD) sequence (Fig. 4C). Translation initiation rates were predicted using the RBS Calculator (22, 23), and SD variants with predicted translation initiation rates approximately 10-fold lower (SDweak) or higher (SDstrong) than that of the native sequence were generated. Under 2.5 mM Zn conditions, the Δ*rpmJ*/WT ratio was approximately fivefold for the wild-type construct but decreased to 1.4-fold for the SDweak construct and to approximately onefold for the SDstrong construct (Fig. 4D, right). These results indicate that enhanced *zntA* expression in the *rpmJ*-deficient strain under excess zinc conditions requires the native *zntA* SD sequence and cannot be explained solely by the predicted translation initiation rate.

### Deletion of *ykgO*, a paralog of *rpmJ*, confers zinc resistance

RpmJ has a paralog, YkgO, whose expression is repressed by Zur under zinc-replete conditions and induced under zinc-deficient conditions (13, 14). Unlike RpmJ, YkgO lacks a zinc-binding motif and replaces RpmJ in the ribosome when expressed (15, 24). We therefore examined whether the zinc-binding motif of RpmJ contributes to zinc resistance by analyzing strains lacking or expressing *ykgO*.

Deletion of *ykgO* alone did not affect zinc resistance (Fig. 5A), consistent with the absence of *ykgO* expression under zinc-replete conditions (14). To induce *ykgO* expression, we deleted the *zur* repressor. The Δ*zur* strain exhibited zinc resistance comparable to that of the wild-type strain (Fig. 5A). However, deletion of *zur* in the Δ*rpmJ* background abolished the zinc resistance conferred by *rpmJ* deficiency (Fig. 5A). Additional deletion of *ykgO* restored zinc resistance in the Δ*rpmJ* Δ*zur* strain (Fig. 5A). These results indicate that induction of *ykgO* expression suppresses the zinc resistance conferred by *rpmJ* deficiency.

**Figure. 5.**
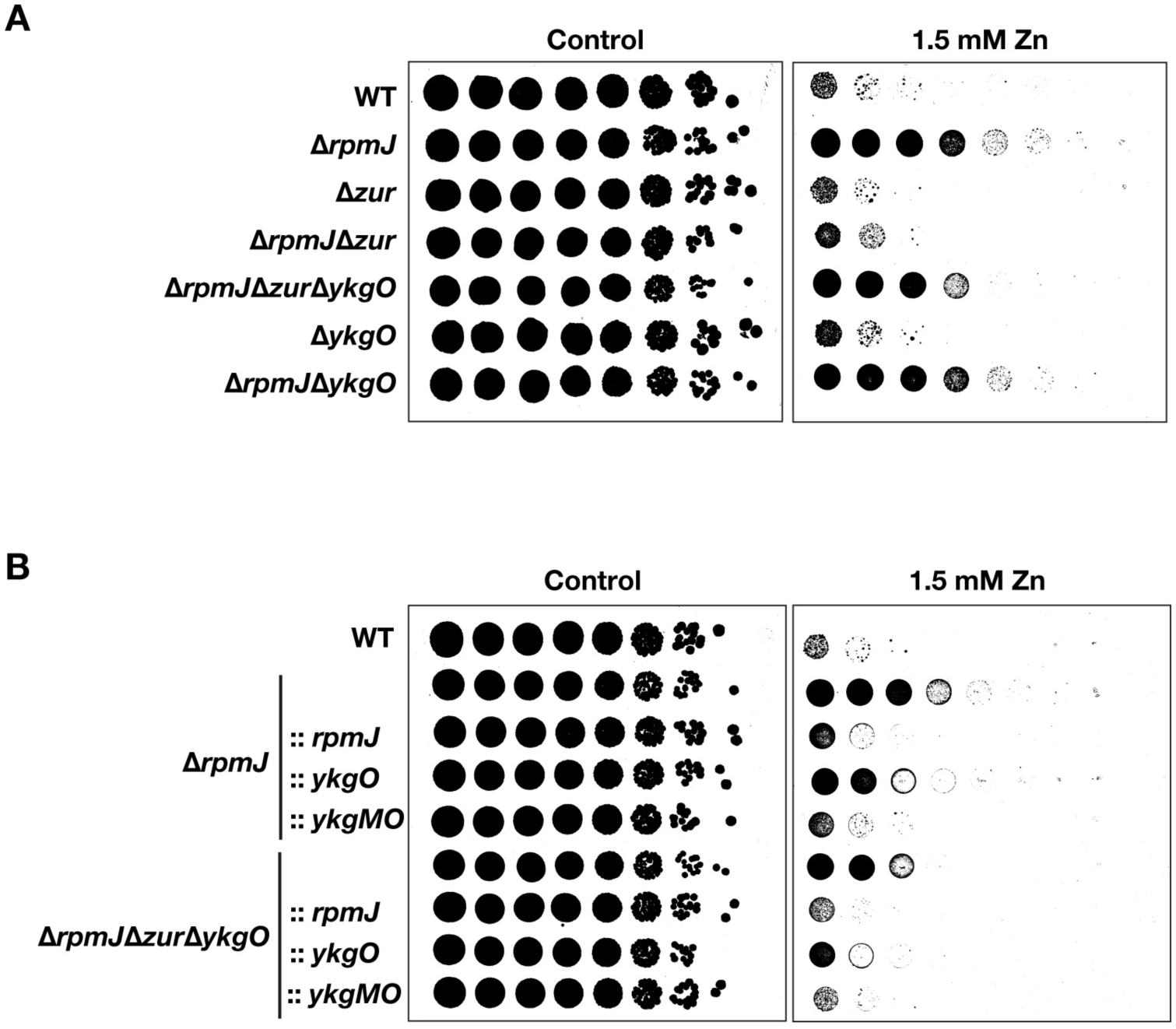
Deletion of *ykgO*, a paralog of *rpmJ*, confers zinc resistance. **(A)** Zinc resistance conferred by *ykgO* deletion. Ten-fold serial dilutions of overnight cultures of the indicated strains were spotted onto LB agar plates containing 1.5 mM Zn or onto LB agar plates without added zinc and incubated for 20 hours. Images are shown after binarization. **(B)** Complementation of *rpmJ* deficiency by *ykgO*. pMW118-rpmJ, pMW118-ykgO, or pMW118-ykgMO was introduced into the Δ*rpmJ* or Δ*rpmJ* Δ*zur* Δ*ykgO* strains. Ten-fold serial dilutions of overnight cultures were spotted onto LB agar plates containing 1.5 mM Zn or onto LB agar plates without added zinc, both supplemented with 1 mM IPTG, and incubated for 20 hours. Images are shown after binarization.

To further examine the role of YkgO, we complemented the Δ*rpmJ* Δ*zur* Δ*ykgO* strain with either *rpmJ*, *ykgO*, or the *ykgMO* operon. Complementation with *rpmJ*, *ykgO*, or the *ykgMO* operon attenuated zinc resistance, with *rpmJ* and the *ykgMO* operon completely abolishing the phenotype and *ykgO* alone producing only a partial reduction (Fig. 5B). Similarly, complementation of the Δ*rpmJ* strain with *ykgO* partially reduced zinc resistance, whereas complementation with the *ykgMO* operon completely abolished it (Fig. 5B). Together, these results indicate that the zinc-binding motif of RpmJ is not required for zinc resistance. Instead, they suggest that altered ribosome composition resulting from the absence of either RpmJ or YkgO promotes zinc resistance.

### *ykgO* deficiency increases *zntA* expression through a mechanism similar to that of

### *rpmJ* deficiency

To investigate the mechanism underlying the zinc resistance conferred by *ykgO* deficiency, we first examined ZntA protein expression. Under 2.5 mM Zn conditions, ZntA protein levels were increased in the Δ*rpmJ* Δ*zur* Δ*ykgO* strain relative to the wild-type strain, whereas no increase was observed in the Δ*rpmJ* Δ*zur* strain (Fig. 6A). This pattern was similar to that observed in the Δ*rpmJ* strain.

**Figure. 6.**
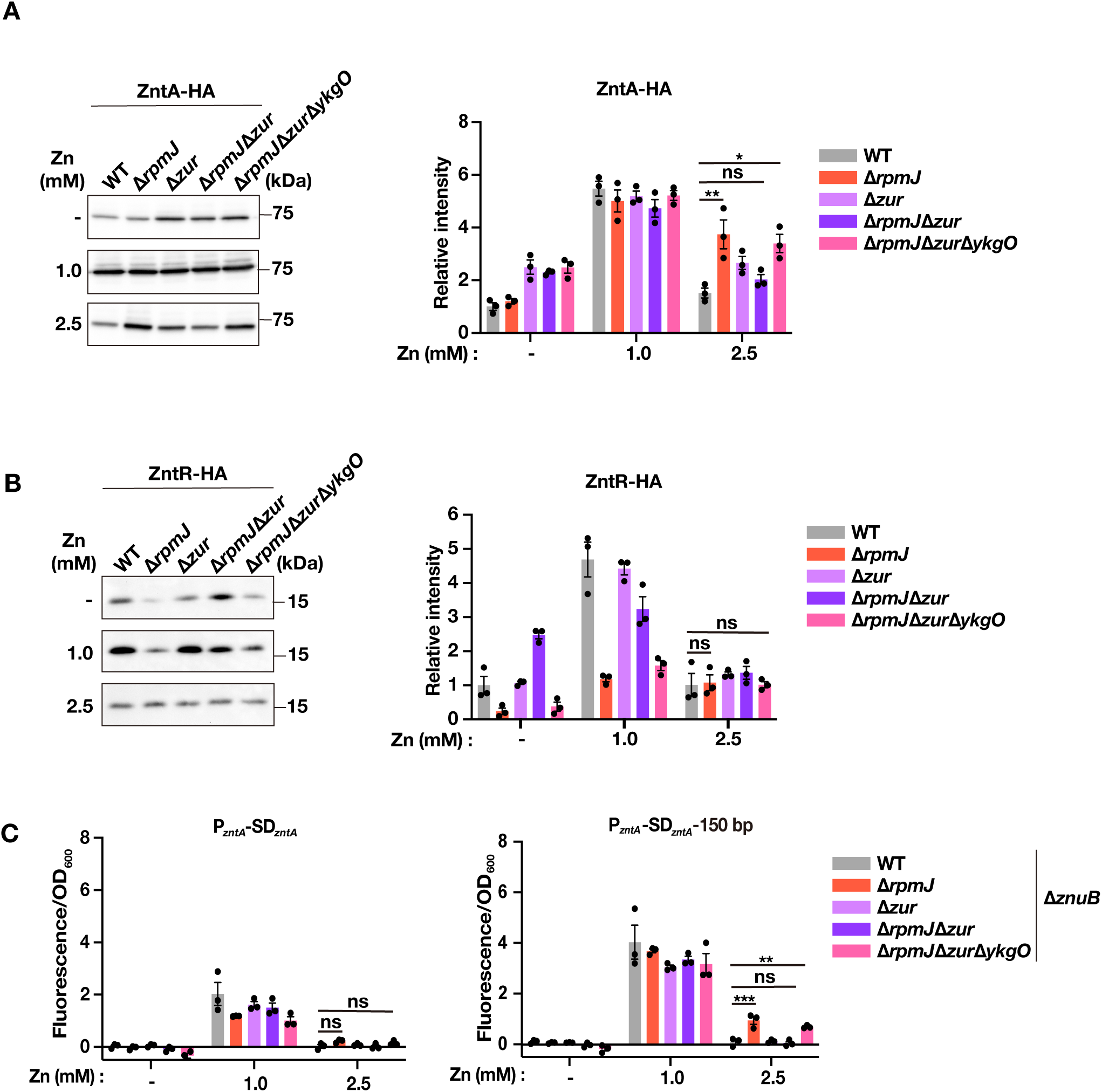
*ykgO* deficiency enhances *zntA* expression through a mechanism similar to that of *rpmJ* deficiency. **(A)** ZntA-HA expression in the Δ*rpmJ* Δ*zur* Δ*ykgO* strain. pMW118-zntA-HA was introduced into the wild-type, Δ*rpmJ*, Δ*zur*, Δ*rpmJ* Δ*zur*, and Δ*rpmJ* Δ*zur* Δ*ykgO* strains. Cells were cultured and analyzed as described in Fig. 2A. Data are presented as mean ± S.E.M. (*n* = 3, \**p* < 0.05, \*\**p* < 0.01; Tukey’s multiple comparisons test). **(B)** ZntR-HA expression in the Δ*rpmJ* Δ*zur* Δ*ykgO* strain. pMW118-zntR-HA was introduced into the wild-type, Δ*rpmJ*, Δ*zur*, Δ*rpmJ* Δ*zur*, and Δ*rpmJ* Δ*zur* Δ*ykgO* strains. Cells were cultured and analyzed as described in Fig. 2A. Data are presented as mean ± S.E.M. (*n* = 3, Tukey’s multiple comparisons test). **(C)** *zntA* reporter activity in the Δ*rpmJ* Δ*zur* Δ*ykgO* strain. Reporter plasmids carrying P*zntA*-SD*zntA*-sfGFP or P*zntA*-SD*zntA*-zntA coding region 150 bp-sfGFP were introduced into the Δ*znuB*, Δ*znuB* Δ*rpmJ*, Δ*znuB* Δ*zur*, Δ*znuB* Δ*rpmJ* Δ*zur*, and Δ*znuB* Δ*rpmJ* Δ*zur* Δ*ykgO* strains. Reporter activity was determined as described in Fig. 3B. Data are presented as mean ± S.E.M. (*n* = 3, \*\**p* < 0.01, \*\*\**p* < 0.001; Tukey’s multiple comparisons test).

We next examined ZntR protein expression. As observed in the Δ*rpmJ* strain, ZntR protein levels were unchanged in the Δ*rpmJ* Δ*zur* Δ*ykgO* strain under 2.5 mM Zn conditions (Fig. 6B).

Finally, we examined whether the *zntA* coding region is required for enhanced *zntA* expression using reporter assays. To avoid the confounding effect of increased zinc uptake caused by derepression of *znuB* in the Δ*zur* background, reporter assays were performed in the Δ*znuB* background. Reporter activity in the Δ*rpmJ* Δ*zur* Δ*ykgO* strain was not increased by the native *zntA* promoter alone but increased when the reporter contained the *zntA* coding region, whereas no increase was observed in the Δ*rpmJ* Δ*zur* strain (Fig. 6C). Together, these results indicate that *ykgO* deficiency enhances *zntA* transcription initiation in a *zntA* coding region-dependent manner through a mechanism similar to that observed for *rpmJ* deficiency.

## DISCUSSION

In this study, we demonstrated that deletion of *rpmJ* enhances *zntA* expression and confers zinc resistance under zinc-excess conditions. Our data further show that enhanced *zntA* expression requires the native *zntA* promoter, the *zntA* coding region, and translation initiation at the native Shine–Dalgarno sequence. Together, these findings indicate that ribosomes lacking RpmJ enhance *zntA* expression through a translation-dependent mechanism.

Our promoter replacement experiments indicate that the increase in *zntA* mRNA observed in the Δ*rpmJ* strain cannot be explained solely by altered *zntA* mRNA stability. Because transcription from the native *zntA* promoter, but not from the constitutive promoter J23106, was enhanced in the Δ*rpmJ* strain (Fig. 3), despite both promoters generating identical *zntA* transcripts, our data are most consistent with increased transcription initiation from the native *zntA* promoter. Furthermore, under 2.5 mM Zn conditions, insertion of the *zntA* coding region specifically enhanced reporter activity in the Δ*rpmJ* strain, with maximal activation observed upon insertion of the first 150 bp of the coding region and little additional increase upon extension to 300 bp (Fig. 3C). This observation suggests that early translation events within the 5′ region of the *zntA* coding sequence are sufficient to promote enhanced transcription. Together with the requirement for the native Shine–Dalgarno sequence, these findings support a model in which recognition of the native Shine–Dalgarno sequence and subsequent translation of the first 150 bp of *zntA* mRNA by ribosomes lacking RpmJ promote transcription initiation from the native *zntA* promoter (Fig. 7, Model ①). To our knowledge, translation has not previously been proposed to stimulate transcription initiation. A recent structural study showed that RNA polymerase (RNAP) can form a complex with the 30S ribosomal subunit, suggesting a mechanism by which RNAP may facilitate formation of the translation initiation complex (25). Although this finding supports communication between the transcription and translation machineries, it does not demonstrate reciprocal regulation whereby translation influences transcription initiation. Moreover, the reported interaction involves the 30S subunit, whereas our results implicate the absence of the 50S protein RpmJ. Whether loss of RpmJ alters ribosome architecture, affects RNAP–ribosome interactions, or influences transcription initiation through another mechanism remains to be determined.

**Figure 7.**
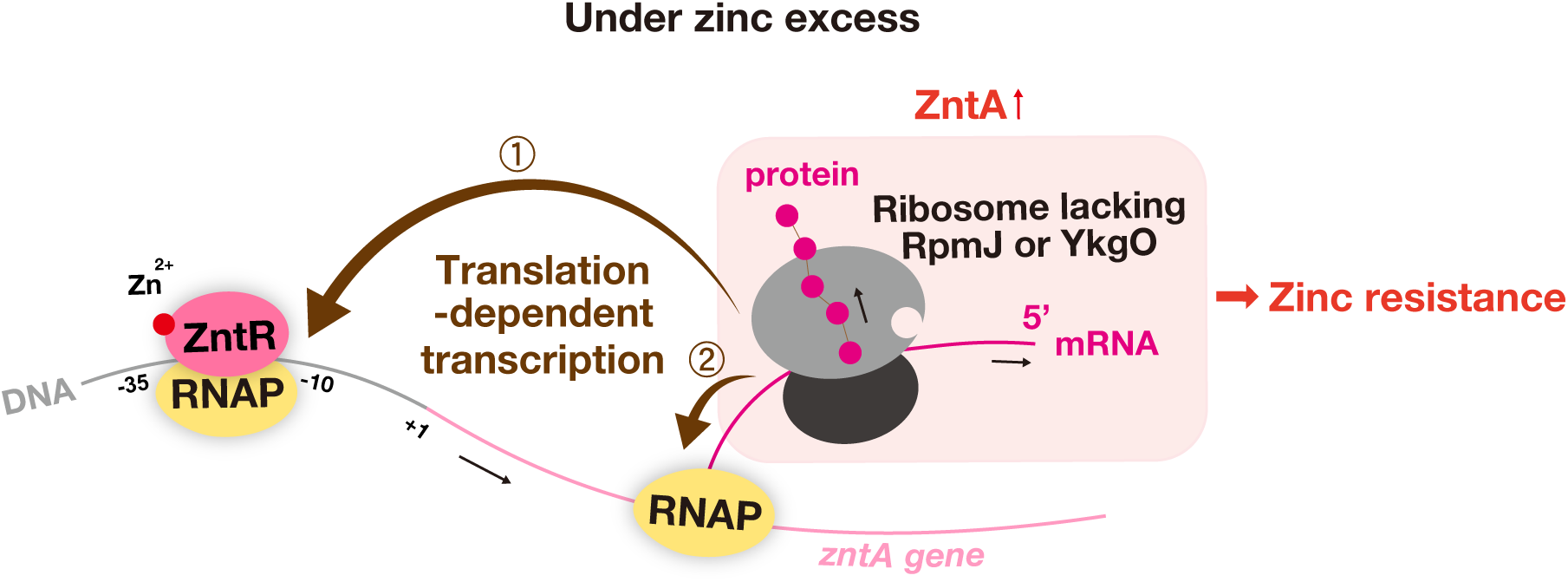
Model for *zntA* upregulation induced by *rpmJ/ykgO* deficiency. Proposed model for the upregulation of *zntA* transcription induced by *rpmJ* or *ykgO* deficiency. Under zinc excess conditions, ribosomes lacking RpmJ or YkgO recognize the native *zntA* Shine–Dalgarno sequence and initiate translation of *zntA* mRNA. In Model ①, this early translation promotes transcription initiation from the native *zntA* promoter, thereby increasing *zntA* mRNA synthesis. In Model ②, the translating ribosome closely follows RNA polymerase (RNAP), thereby promoting transcription continuation and increasing *zntA* expression.

An alternative, and not mutually exclusive, explanation is that ribosomes lacking RpmJ promote transcription elongation (Fig. 7, Model ②). Translation and transcription are tightly coupled in bacteria, and actively translating ribosomes suppress Rho-dependent premature transcription termination by remaining closely associated with RNAP (26, 27). Ribosomes also prevent RNAP backtracking, thereby promoting transcription elongation (28). More recently, rapid translation initiation and efficient translation have been shown to contribute to sustained transcription and maintenance of transcription–translation coupling (29, 30). Consistent with these observations, ribosomes lacking RpmJ may recognize the native *zntA* Shine–Dalgarno sequence more efficiently, thereby maintaining close coupling with RNAP and suppressing premature transcription termination. However, because the increase in *zntA* expression caused by *rpmJ* deficiency required the native *zntA* promoter, this mechanism alone is unlikely to fully account for our observations. Future studies will be required to determine the relative contributions of Models ① and ②.

A second important finding of this study is that the zinc resistance conferred by *rpmJ* deficiency was abolished by expression of its paralog *ykgO*. Because YkgO lacks the zinc-binding motif present in RpmJ, these results argue against a model in which zinc released from RpmJ directly induces *zntA* expression. Instead, our data indicate that altered ribosome composition resulting from the absence of either RpmJ or YkgO is responsible for this phenotype. Consistent with this interpretation, *ykgO* deficiency increased *zntA* expression through a mechanism similar to that observed in the Δ*rpmJ* strain. The RpmJ–YkgO exchange system has been proposed to function as a zinc reservoir by releasing zinc from RpmJ under zinc-deficient conditions (31, 32). Our findings reveal an additional role for this system by linking ribosome composition to regulation of *zntA* expression and zinc resistance.

Ribosome heterogeneity has emerged as an important mechanism for selective translation in both bacteria and eukaryotes(33, 34). In bacteria, kasugamycin induces formation of 61S ribosomes lacking several ribosomal proteins, which preferentially translate leaderless mRNAs (35). Likewise, MazF-dependent processing of 16S rRNA has been proposed to generate specialized ribosomes that selectively translate leaderless transcripts (36), although the physiological relevance of this mechanism remains controversial (37). In eukaryotes, environmental stress reduces incorporation of Rps26 into ribosomes, thereby altering translation of stress-responsive mRNAs according to Kozak sequence strength and contributing to stress adaptation (34). Our findings extend this concept by suggesting that changes in ribosome composition can regulate bacterial stress resistance not only through selective translation but also by coupling translation of specific mRNAs to transcriptional regulation.

In summary, our study identifies a previously unrecognized mechanism linking ribosome composition to bacterial zinc resistance. Ribosomes lacking RpmJ enhance *zntA* expression through a translation-dependent mechanism that requires the native *zntA* promoter, coding region, and Shine–Dalgarno sequence. Future biochemical reconstitution experiments using purified ribosomes lacking RpmJ will be required to elucidate the molecular mechanism by which these ribosomes directly promote transcription initiation from the native *zntA* promoter.

## MATERIALS AND METHODS

### Bacterial strains and culture conditions

*Escherichia coli* BW25113 and its derivative strains were routinely cultured on LB agar plates or in LB broth at 37°C with aeration. Strains carrying pMW118 or its derivatives were cultured in LB medium supplemented with ampicillin (100 µg/ml). The bacterial strains and plasmids used in this study are listed in Tables 1 and 2. For gene expression analyses, overnight cultures were diluted 1:100 into fresh LB medium and incubated for 2 hours. Samples collected at this time point served as the samples without added zinc. For zinc-treated samples, zinc was added to final concentrations of 0.5–2.5 mM after 2 hours of growth, and cultures were incubated for an additional 1 hour before sample collection.

**Table 1.**
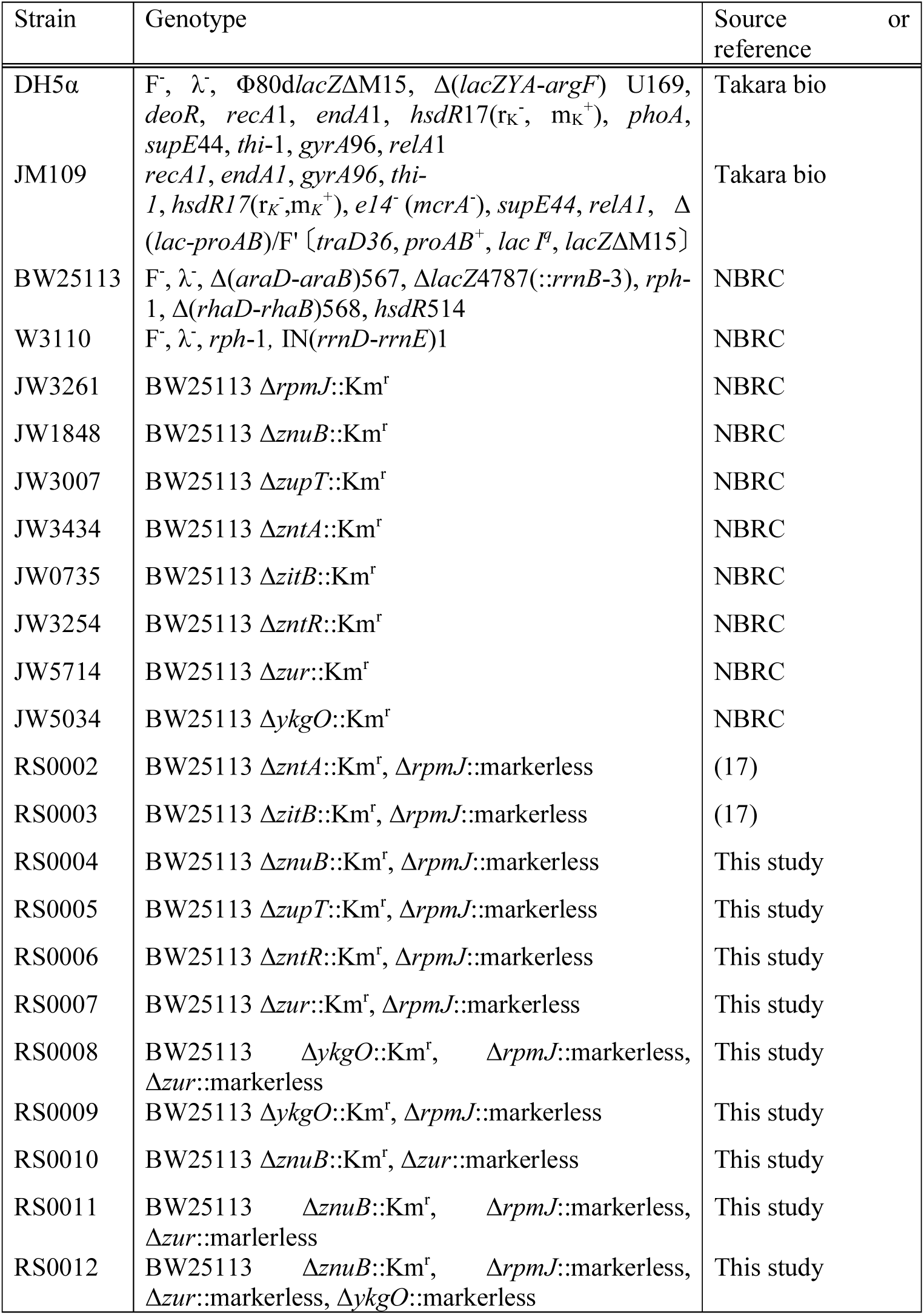
Bacterial strains used in this study.

**Table 2.**
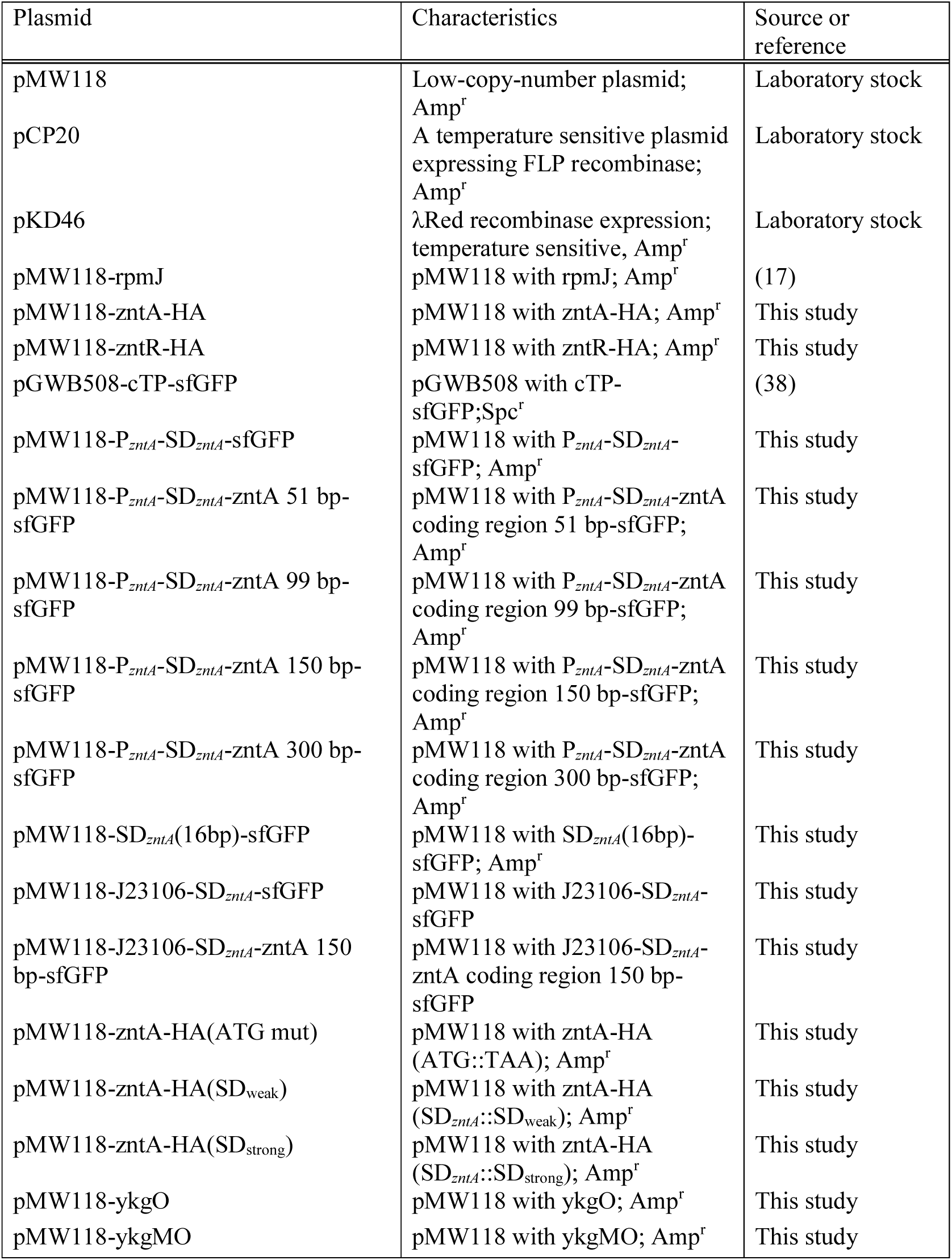
Plasmids used in this study.

### Construction of gene knockout mutants

Single-gene deletion strains were generated by P1 phage transduction of the corresponding mutations from the Keio collection into *E. coli* BW25113. Multiple-deletion strains were constructed by repeated cycles of P1 phage transduction followed by removal of the kanamycin resistance (Kmr) cassette using the FLP–FRT recombination system. Briefly, the Kmr cassette was excised by introducing pCP20, which expresses FLP recombinase, and the resulting markerless strain was used as the recipient for subsequent rounds of P1 transduction. The Δ*rpmJ* Δ*zntR* strain was constructed by One-Step Inactivation. A PCR product containing the *zntR::*FRT*-*KmR-FRT allele together with approximately 500 bp of upstream and downstream homologous sequences was amplified from the Δ*zntR* strain and introduced into the markerless Δ*rpmJ* strain carrying pKD46. All gene deletions were confirmed by PCR.

### Evaluation of bacterial resistance to zinc

LB agar was autoclaved, supplemented with ZnSO4·7H2O (Nacalai Tesque), and poured into square Petri dishes (Eiken Chemical). Overnight cultures of *E. coli* were serially diluted 10-fold in 96-well plates, and 5 µl of each dilution was spotted onto LB agar plates containing zinc. For complementation experiments, IPTG was added to a final concentration of 1 mM. The plates were incubated at 37°C for 20 hours, and colony growth was photographed.

### Cloning of plasmid constructs

To construct plasmids expressing HA-tagged ZntA or ZntR, the *zntA* and *zntR* coding regions were amplified from the *Escherichia coli* BW25113 genome by PCR using primers encoding a C-terminal HA tag and cloned into the XbaI and HindIII sites of pMW118 to generate pMW118-zntA-HA and pMW118-zntR-HA, respectively. pMW118-zntA-HA (ATG mut) was generated by PCR-based site-directed mutagenesis using the primers zntA ATG mutation_F and zntA ATG mutation_R. pMW118-zntA-HA (SDweak) and pMW118-zntA-HA (SDstrong) were generated by inverse PCR using pMW118-zntA-HA as the template, followed by phosphorylation with T4 Polynucleotide Kinase (Takara) and self-ligation. The *ykgO* gene and the *ykgMO* operon were amplified from the BW25113 genome and cloned into the EcoRI and BamHI sites of pMW118 to generate pMW118-ykgO and pMW118-ykgMO, respectively.

To construct the reporter plasmids, pMW118-P*zntA*-SD*zntA*-lacZ was first generated. The *lacZ* gene was amplified from the *E. coli* W3110 genome and cloned into the XbaI and HindIII sites of pMW118. The *zntA* promoter and Shine–Dalgarno sequence were then amplified from BW25113 genomic DNA and inserted into the KpnI and BamHI sites upstream of *lacZ*. The *sfGFP* coding region was amplified from pGWB602-cTP-sfGFP (Ishikawa *et al*., 2023) and assembled with linearized pMW118-P*zntA*-SD*zntA*-lacZ using the NEBuilder HiFi DNA Assembly Master Mix (New England Biolabs) to generate pMW118-P*zntA*-SD*zntA*-sfGFP. pMW118-SD*zntA* (16 bp)-sfGFP was generated by inverse PCR using pMW118-P*zntA*-SD*zntA*-sfGFP as the template.

For reporter constructs containing the *zntA* coding region, the full-length *zntA* coding region amplified from BW25113 genomic DNA was inserted into linearized pMW118-P*zntA*-SD*zntA*-sfGFP using the SLiCE method to generate pMW118-P*zntA*-SD*zntA*-zntA coding region-sfGFP. Reporter constructs containing the first 51, 99, 150, or 300 bp of the *zntA* coding region were generated from this plasmid by inverse PCR followed by SLiCE-mediated ligation.

The J23106 promoter (Registry of Standard Biological Parts, http://parts.igem.org/Part:BBa_J23106) constructs were generated from pMW118-P*zntA*-SD*zntA*-sfGFP or pMW118-P*zntA*-SD*zntA*-zntA150 bp-sfGFP by replacing the native *zntA* promoter with the constitutive promoter J23106 using inverse PCR followed by SLiCE-mediated ligation.

### Western blot analysis

Cells corresponding to an OD600 of 0.5 were harvested (1 ml), resuspended in TE buffer and 3× SDS sample buffer, and boiled for 5 min. Proteins were separated on 12.5% or 15% SDS–PAGE gels and transferred onto an Immobilon-P PVDF membrane (Millipore). Membranes were blocked with TBS-T (20 mM Tris-HCl, pH 7.4, 150 mM NaCl, 0.1% Tween-20) containing 5% (w/v) skim milk overnight at 4°C or for 30 min at room temperature. The membranes were incubated with anti-HA tag antibody (MBL, clone TANA2) diluted 1:10,000 in Can Get Signal Solution 1 (TOYOBO) for 40 min at 37°C, followed by incubation with HRP-conjugated anti-mouse IgG antibody (Promega, W402B) diluted 1:25,000 in Can Get Signal Solution 2 (TOYOBO) for 40 min at 37°C. Signals were detected using an iBright 1500 imaging system (Thermo Fisher Scientific) and quantified with ImageJ.

### RNA extraction and RT-qPCR

Collected cells were mixed with water-saturated phenol and immediately frozen in liquid nitrogen. Samples were resuspended in TE buffer containing SDS and lysozyme and incubated at 65°C for 2 min. Total RNA was extracted using the RNeasy Mini Kit (Qiagen). RNA concentrations were determined using a NanoDrop One spectrophotometer (Thermo Fisher Scientific), and equal amounts of RNA were used for subsequent analyses. cDNA was synthesized using MultiScribe Reverse Transcriptase and random hexamer primers (Invitrogen, Thermo Fisher Scientific). Quantitative PCR was performed using KOD SYBR qPCR Mix (TOYOBO) on a StepOnePlus Real-Time PCR System (Thermo Fisher Scientific). Transcript levels were normalized to *gyrA*. The primers used in this study are listed in Table 3.

**Table 3.**
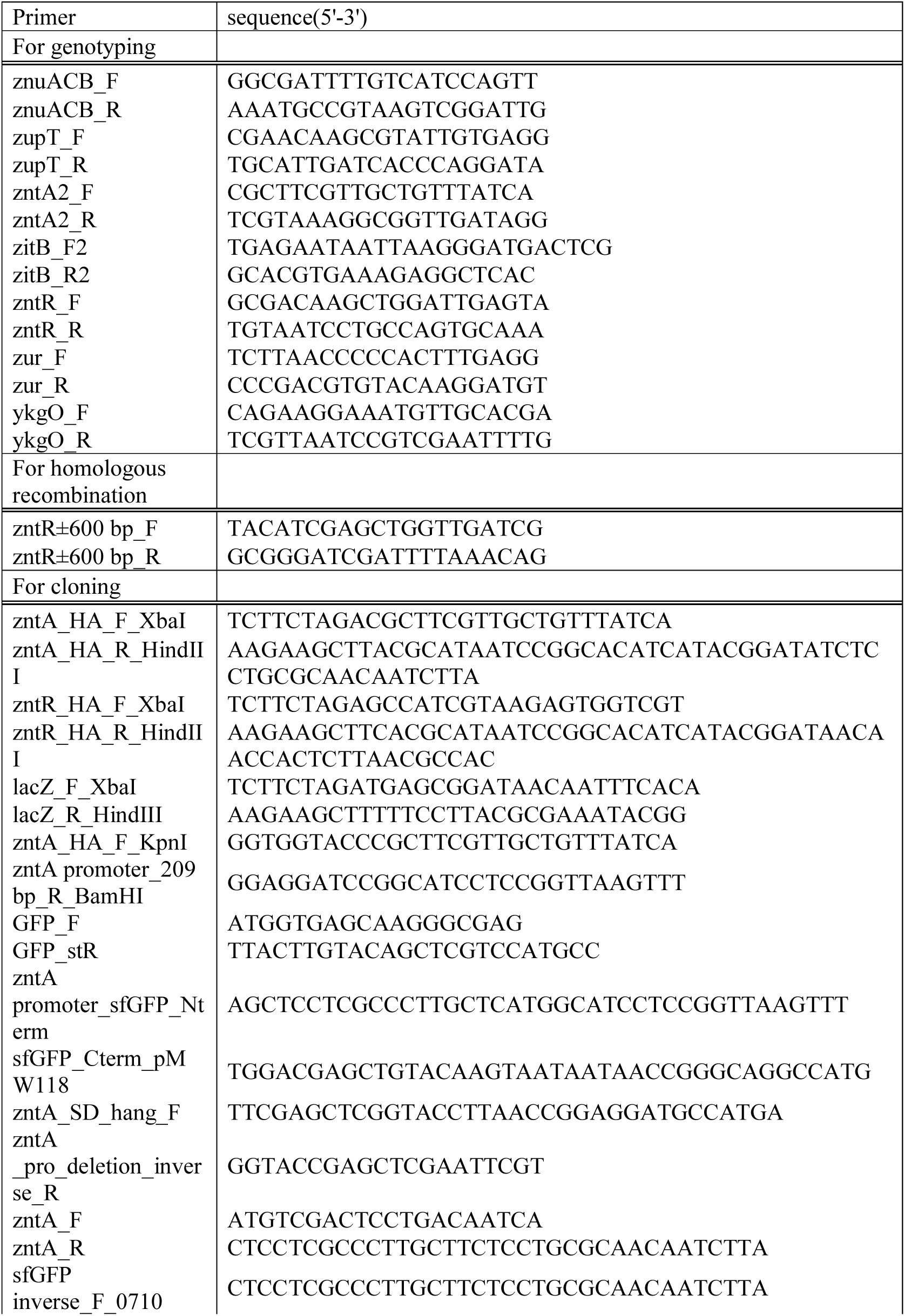
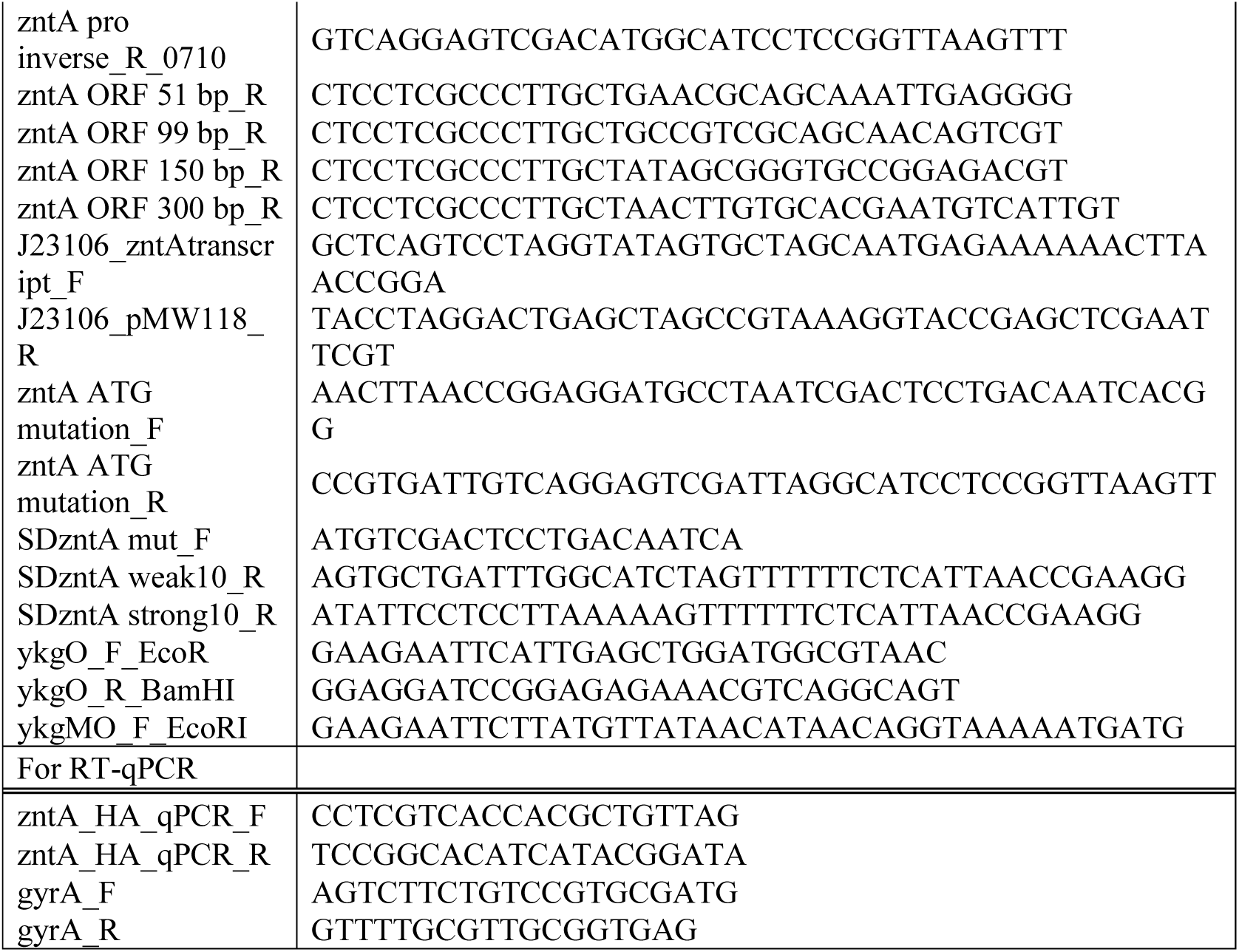
Oligonucleotide primers used in this study.

### Reporter assay

At the time of sampling, the OD600 of each culture was measured. Equal volumes of culture were harvested by centrifugation, and the cell pellets were resuspended in saline and transferred to a 96-well plate. Fluorescence was measured using a Fluoroskan Ascent CF microplate fluorometer (Labsystems; currently Thermo Fisher Scientific) with excitation at 485 nm and emission at 527 nm.

Reporter activity was calculated as fluorescence intensity normalized to OD600. Background reporter activity, determined using cells carrying pMW118-SD*zntA* (16 bp)-sfGFP as the promoterless construct, was subtracted from all values.

### Design of Shine–Dalgarno sequence variants

Shine–Dalgarno sequence variants were designed using the RBS Calculator (https://salislab.net/software/login). The predicted translation initiation rate of the native *zntA* Shine–Dalgarno sequence was first calculated, and variant sequences with predicted translation initiation rates approximately 10-fold lower or higher than that of the native sequence were subsequently designed.

## ACKNOWLEDGMENTS

We thank the National BioResource Project-*E. coli* (National Institute of Genetics, Japan) for providing the Keio collection. This work has been partly supported by the Core-Facility Portal (CFPOU) at Okayama University (DGP_508, DGP_653). This study was supported by Japan Society for the Promotion of Science (JSPS) Grants-in-Aid for Scientific Research (Grants 23K24131, 23K06130, 24K01760, 26K02250, 26K23680, 24K21872) and the JSPS research fellowships for young scientists (24KJ1719 to RS).

